# Acute drug induced parkinsonian akinesia is associated with reduced rate and increased regularity of neuronal activity in the pedunculopontine tegmental nucleus of unanesthetized rats

**DOI:** 10.1101/2024.05.14.592858

**Authors:** Xiaodong Lu, Jeffery R. Wickens, Brian I. Hyland

## Abstract

The pedunculopontine tegmental nucleus of the brainstem is important for a wide range of functions, including sensorimotor integration and regulation of locomotion and discrete forelimb movements. Evidence suggests it is structurally disordered as part of the neuropathology of the movement disorder Parkinson’s disease. To assess whether neuronal activity is also affected in parkinsonian model animals we used single neuron recordings in free moving rats to determine the effects of acute drug-induced parkinsonism on the firing rate of the neurons. Acute parkinsonian akinesia was associated with reduction in resting firing rate and increased firing regularity, consistent with reduced excitability in this state.

## Introduction

The pedunculopontine tegmental nucleus of the brainstem (PPTg) is implicated in multiple aspects of behavioural regulation, including the control of movement. Underpinning its motor control functions are afferent inputs from motor control structures including direct projections from cortical motor regions (Matsumura *et al*., 2000), and onward projections to brainstem and spinal cord motor centres (Inglis & Winn, 1995; Winn *et al*., 1997). The PPTg is a major component of the “mesencephalic locomotor region”, involved in the generation and pacing of stepping movements (Garcia-Rill *et al*., 1983; Garcia-Rill *et al*., 1987; Garcia-Rill & Skinner, 1988; Garcia-Rill *et al*., 1990; Karachi *et al*., 2010; Roseberry *et al*., 2016; Caggiano *et al*., 2018). Increasing evidence also implicates it in the regulation of discrete skilled forelimb movements (Dunbar *et al*., 1992; Matsumura *et al*., 1997; Dormont *et al*., 1998; Florio *et al*., 1999; Weinberger *et al*., 2008; MacLaren *et al*., 2014; Lau *et al*., 2015; Lu *et al*., 2024).

Consistent with a potential role in normal movement control, several lines of evidence suggest that changes in PPTg activity may be important in the pathogenesis of movement disorders in Parkinson’s disease (Pahapill & Lozano, 2000; Tubert *et al*., 2019). Lesions of PPTg in monkeys produce bradykinesia, similar to mild Parkinsonism (Kojima *et al*., 1997; Aziz *et al*., 1998; Munro-Davies *et al*., 1999; Takada *et al*., 2000). Conversely, deep brain stimulation of the nucleus has been reported to ameliorate parkinsonism in both primate models (Jenkinson *et al*., 2004; Jenkinson *et al*., 2006) and in Parkinson’s disease patients (Thevathasan *et al*., 2018). However, the correlates of parkinsonism at the level of neuronal activity in PPTg remain unclear due to conflicting results in the existing literature. Thus, using various measures of level of activation of PPTg neurons, some studies report no change (Heise & Mitrofanis, 2006; Aravamuthan *et al*., 2008), others found increases (Carlson *et al*., 1999; Orieux *et al*., 2000; Breit *et al*., 2001; Jeon *et al*., 2003; Zhang *et al*., 2008; Geng *et al*., 2016), while still others decreases in the measured parameter (Mitchell *et al*., 1989; Gomez-Gallego *et al*., 2007). Here, we examined the effects of acute parkinsonism induced by the dopamine antagonist raclopride on the level of resting activity of the neurons. The results indicated that acute parkinsonism induced by the dopamine antagonist raclopride is associated with reduction in average neuronal resting firing rate, and increased firing regularity.

## Methods

All procedures were approved by the University of Otago Committee on Ethics in Care and Use of Laboratory Animals and were in accord with the “Principles of Laboratory Animal Care” (NIH publication number 80-23, revised 1996). Rats were maintained on a reverse day-night cycle, and all handling and experimental sessions were conducted in a darkened room under red light during the animal’s dark phase.

### Animals and surgery

Male Wistar rats (∼ 400 g at the time of surgery) were habituated over two weeks to handling and to being placed in a clear acrylic task box, and then underwent surgery to implant chronic extracellular recording electrodes into the contralateral PPTg (anteroposterior (AP) -7.8 mm and lateral (L) ± 2.0 mm, relative to bregma (Paxinos & Watson, 1997)) using sterile stereotaxic technique under full anaesthesia (pentobarbitone 60 mg/kg i.p.). The tips of recording electrodes were left just dorsal to the PPTg at 6.5 mm from the dura. Recording electrodes were eight 0.007” formvar-coated nichrome microwires (A-M Systems Inc., Carlsborg, Washington, U.S.A), mounted in a “Scribe” on-head microdrive (Bilkey & Muir, 1999), to allow gradual advance of the electrodes through the PPTg over 1-3 months. The microdrive assembly was fixed in place with skull screws and dental acrylic that also anchored the connection plug (MacIntyre connector, Ginder Scientific).

### Experimental procedures

During recording sessions, signals from electrodes were amplified and filtered (2000 -8000 x, 0.5 - 10 kHz bandpass). Extracellular action potentials from single neurons were recorded, along with marker channels for automatically or manually entered behavioural events using SciWorks software (Datawave, CO). These rats were also trained in a forelimb reaching task and neurons tested for responses during reaching and to an auditory stimulus; results from that part of the study have been published elsewhere (Lu *et al*., 2024). Approximately 10 minutes following completion of the other experiments the effect of neuroleptic-induced akinesia (catalepsy) on baseline firing of PPTg neurons was assessed. At this stage the animals were satiated and well habituated to the environment, and so resting quietly in the test box with little spontaneous movement. Cell activity was recorded for 5 minutes to establish the activity level during natural quiet rest. Animals were then injected with a cataleptic dose of the selective dopamine D2 antagonist raclopride (1.5 mg/kg in saline i.p.), and 20 minutes later another 5-minute block of data was recorded (Fig. 1 A). During neural recording periods animal behaviour was monitored and locomotor, grooming or rearing episodes marked on the data file. The grid test (Morelli & Di Chiara, 1985) was used to confirm that the drug had been effectively administered. For this, animals were placed on a vertical wire grid and the time that elapsed until movement of one paw to a new location recorded. All neurons reported here were recorded in sessions where the grid time was > 60 s indicating strong catalepsy, compared to < 5 s in all pre-drug tests.

**Figure 1.**
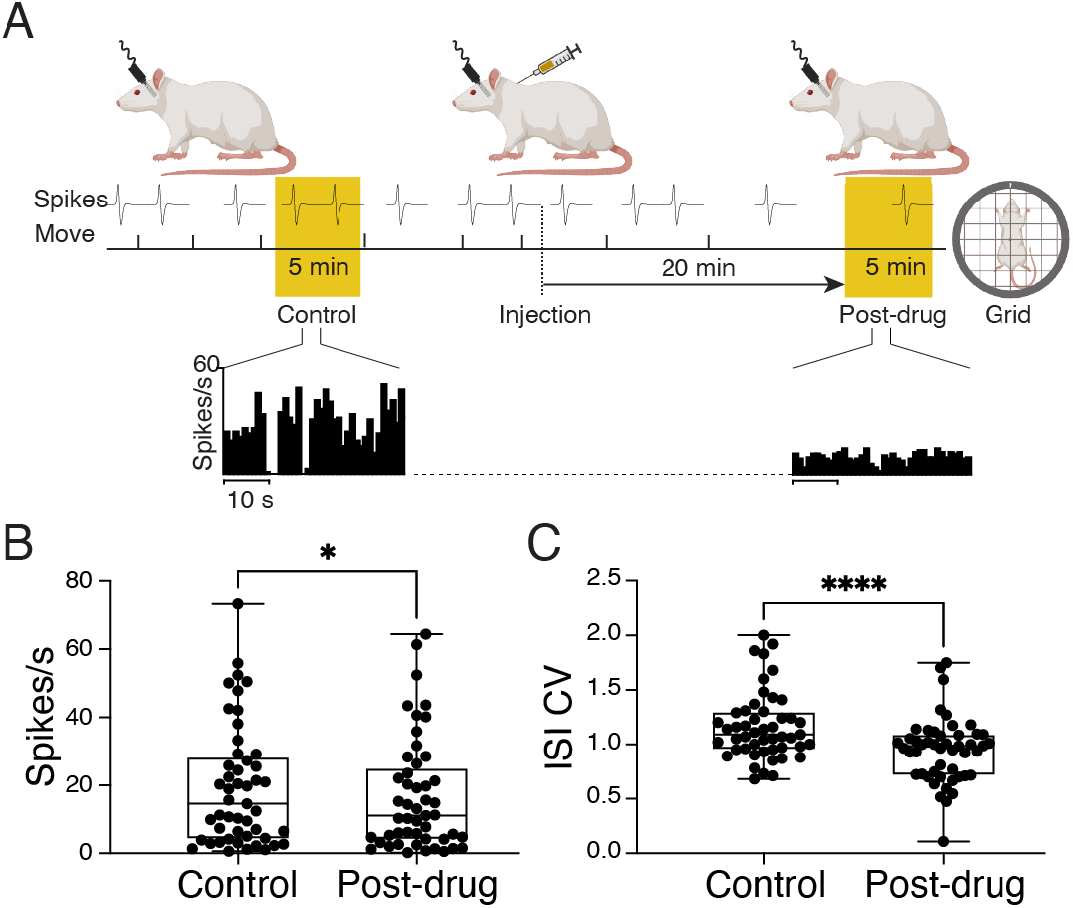
**A**. Top part of the panel is a schematic showing recording protocol. Rats implanted with chronic indwelling recording electrodes rested quietly in a task box while action potentials (Spikes) from PPTg neurons were recorded, and movement episodes were manually entered on a marker channel (Move). Following the recordings, the grid test was performed to confirm catalepsy. Rate-meter histograms display segments of actual data from an example recorded neuron, converted to instantaneous firing rate in 1 s bins. Time scale bar = 10 s. **B**. Graph shows all points data for firing rate. Box indicated 25^th^, 75^th^ percentiles and whiskers show minimum and maximum. *, p < 0.05, paired t-test. **C**. Graph shows all points and box and whiskers plot for coefficient of variation of interspike intervals (ISI CV). ****, p < 0.0001, paired t-test/s

On completion of recordings, lesions were induced at the tips of selected electrodes by passing a DC current of 2 mA for at least 15 seconds, and one week later the animals were sacrificed with an anaesthetic overdose and the brains perfused and processed for histological reconstruction of the recording bundle placement.

### Data analysis

Unit activity was analysed off-line using Spike2 software (CED, Cambridge, UK). Spikes belonging to single neurons were discriminated based on waveform shape. For analysis of neural activity before and after drug administration, mean firing rate was calculated from the rate-meter histogram of each spike train over 5 min (1 s bins). The coefficient of variation of interspike intervals (CV), a measure of firing variability, was calculated from the same time periods using interspike interval histograms with 10 ms bins and including intervals up to 1 s. Animals were uniformly cataleptic throughout the post drug period. To reduce differences between pre- and post catalepsy that could be caused by differences in the overall level of physical activity, for the control data we selected epochs from the pre-drug control recordings that showed low levels of spontaneous movement, using the behaviour markers entered during recording.

Quantitative averaged data are presented in text as mean ± 1 standard deviation. Statistical analyses were performed using GraphPad Prism Version 8 (GraphPad Software LLC).

## Results

A total of 49 isolated single neurons were recorded throughout a 5-minute pre-injection control period of quiet rest, and a further 5-minute post-injection period during catalepsy induced by injection of the dopamine D2-receptor antagonist raclopride. Grid tests confirmed successful catalepsy induction in all cases, with rats moving a paw in less than 5 s in all pre-drug control tests whereas no rats re-positioned a paw within 60 seconds in the post drug tests. Matching this, during post drug recordings, there was profound catalepsy, with no movements occurring once recording started in many cases. Overall, during the total 5 minutes pre-drug recording, quiet rest was confirmed by very low occurrence of movement, with markers occurring at a mean rate of 0.121 ± 0.041 events/s. To further match movement activity across pre and post-drug periods, for the control we used a period of minimal movement events from within the pre-drug period as baseline for analysis. As intended, activity during this period even more closely approached zero (0.056 ± 0.035 events/s, t_(35)_ = 10.19, p < 0.001 paired t-test compared to the whole file).

Neural activity during the control and cataleptic periods was characterised by calculating mean firing rate, and the coefficient of variation of interspike intervals (CV) as a measure of regularity of cell firing. Example rate-meter histogram data are illustrated in Fig. 1 A, showing a clear decline in firing rate in this neuron during drug induced catalepsy compared to pre-injection quiet rest. This was confirmed at the population level by the firing rate analysis shown in Fig. 1 B. During quiet rest prior to drug injection, tested neurons had a mean firing rate of 19.42 ± 17.71 spikes/s. During drug induced catalepsy, there was a significant reduction in baseline firing rate, with a mean of 16.75 ± 16.6 spikes/s (t_(48)_ = 2.018, p < 0.05, paired t-test).

The analysis for CV of interspike intervals is shown in Fig. 1 C. This measure prior to drug administration averaged 1.162 ± 0.31 across all neurons. Following induction of catalepsy under dopamine D2 receptor blockade it was significantly reduced to 0.946 ± 0.299 (t_(48)_ = 4.576, p < 0.0001). This finding indicates a reduced variability, i.e. increased regularity, of spike firing pattern under the influence of the drug.

The tested neurons were also recorded during performance of a reach to grasp task, and during exposure to an auditory stimulus, with these data published previously (Lu *et al*., 2024). We examined whether effects of raclopride on firing rate and CV may differ depending on the activity of neurons during the reach task or in response to tone. For reach relationships, neurons were classified as showing increased firing rate (+) decreased rate (-) or no change (0) during the reach. Analysis with 2-way ANOVA using factors *Time* (before drug, after drug) and *Group* ((+), (-), (0)), there were no significant differential effects, with non-significant *Group* x *Time* interactions for both firing rate (F_(2, 46)_ = 0.26, P = 0.77), and CV (F_(2, 46)_ = 0.19, P = 0.83). Similarly, for neurons split by whether they responded to tone (2 way ANOVA with Factors *Time* (before drug, after drug) and *Tone response* (yes, no)), there was again no significant interaction for either measure (F_(1, 34)_ = 0.63, P = 0.43; F_(1, 34)_ = 1.12, P = 0.3). Thus, drug effects appear independent of these other properties.

## Discussion

Under conditions of drug induced acute parkinsonism we found a small but significant fall in baseline firing rate, accompanied by an increase in regularity as measured by the interspike interval coefficient of variability.

In the light of its likely roles in movement control, it is not surprising that changes in PPTg activity may also be important in the pathogenesis of movement disorders in Parkinson’s disease (Pahapill & Lozano, 2000; Tubert *et al*., 2019). Such acute reduction in firing rate is consistent with previous findings of reduced firing rate in a chronic model of parkinsonism (Florio *et al*., 2007). Parkinsonism is associated with increased activity in inhibitory inputs to PPTg, providing a mechanism for such reductions (Mitchell *et al*., 1989). Further, blockade of inhibitory transmission in PPTg can ameliorate akinesia in parkinsonian monkeys (Nandi *et al*., 2002).

However, there are contradictory findings regarding the neural correlates of parkinsonism in the PPTg, with other single neuron recording studies finding no change (Aravamuthan *et al*., 2008) or increases (Breit *et al*., 2001; Jeon *et al*., 2003; Zhang *et al*., 2008; Geng *et al*., 2016) in baseline firing rate. There is no obvious consistent difference such as the model used or state of anaesthesia that could potentially account for these differences between studies. In all cases the absolute amplitudes of the firing rate changes are small and so could be susceptible to subtle differences in methodology.

Overall, changes in baseline firing rate may provide some window on PPTg neuronal excitability resulting from altered direct dopamine innervation to PPTg (Ryczko & Dubuc, 2017), or on changes in wider basal ganglia and motor cortex circuit interactions that could lead to reduced net tonic excitatory afferent drive at rest. Such reduced activity in the nucleus fits with classical hypotheses about changes in activity in basal ganglia circuits in Parkinson’s disease (Bezard & Gross, 1998; Pahapill & Lozano, 2000). Whatever underlying mechanism is at work, changes at rest could reflect processes that would also come into play when attempts are made to move, and so reflect underlying factors contributing to parkinsonian brady- and akinesia. Further work is required to determine the underlying mechanisms of the subtle changes in resting firing rate and pattern in the parkinsonian PPTg, and whether these mechanisms have functional impact on the ability to initiate or achieve normal speeds of movement (Boraud *et al*., 2002).

## Acknowledgements

Supported by grants from the Neurological Foundation of New Zealand and the NZ Lottery Grants Board

## Data sharing statement

The data that support the findings of this study are available from the corresponding author upon reasonable request.

